# Small molecule inhibitors of G9a reactivate the maternal PWS genes in Prader-Willi-Syndrome patient derived neural stem cells and differentiated neurons

**DOI:** 10.1101/640938

**Authors:** Hao Wu, Carrie Ng, Vivian Villegas, Stormy Chamberlain, Angela Cacace, Owen Wallace

**Affiliations:** Fulcrum Therapeutics, 5th Floor, 26 Landsdowne st., Cambridge, MA, 02142; Genetics and Genome Sciences, University of Connecticut Health Center, 400 Farmington Avenue, Farmington, CT 06030-6403

**Keywords:** Prader-Willi-Syndrome, imprinting, reactivation, G9a, NanoString, 5-Aza, neuron, inhibitor

## Abstract

Patients with Prader-Willi-Syndrome (PWS) display intellectual impairment, hyperphagia, and various behavioral problems during childhood that converge on a neurologic deficit. The majority of PWS patients have genetic deletions of the paternal 15q11–q13 chromosomal region, with their maternal PWS locus intact but epigenetically silenced by hypermethylation and repressive histone modulation of the PWS imprinting center (PWS-IC). Inhibition of the euchromatin histone methyltransferase G9a by small molecules has been recently reported to reactivate PWS genes in patient fibroblasts and a mouse model. However, it is unknown if inhibition of G9a could have similar effect in human PWS neural cells, the cell types that have direct pathophysiological relevance to PWS. Here, we use neural progenitor cells (NPCs) and cortical excitatory neurons derived from a patient iPSC to model PWS, and quantitatively profile the expression of PWS genes using a NanoString panel. We demonstrated that the methylation of the PWS-IC is stable during neuronal lineage conversion, and that the maternal PWS genes remain silenced in PWS NPCs and neurons. Multiple small molecule inhibitors of G9a activate maternal PWS genes in a dose dependent manner in both NPCs and neurons. In addition, G9a inhibitors induce *GNRH1* and *HTR2C*, two neuronal specific genes that contribute to PWS pathology in neurons. Interestingly, distinct from 5-Azacytidine, G9a inhibition does not induce methylation changes of the maternal PWS-IC, indicating that disruption of the histone repressive complex alone is sufficient to drive an open chromatin state at the PWS-IC that leads to partial reactivation of PWS genes.

**Highlights:**

- Modeling PWS disease in a dish using patient derived NPCs and neurons
- G9a inhibition activates maternal PWS genes in patient-derived neural cells
- G9a inhibition activates maternal *SNORD116* and other PWS genes in patient-derived neurons
- Inhibition of G9a induces PWS downstream genes *GNRH1* and *HTR2C* in PWS neurons

## Introduction

Prader–Willi syndrome (PWS) is a developmental disorder in which boys and girls present with developmental delays, including muscle hypotonia, short stature at infancy, intellectual impairment and various behavioral problems during childhood. Most PWS patients exhibit severe hyperphagia and lose appetite control, leading to obesity and type II diabetes (Angulo et al., 2015; Cassidy et al., 2000) when diet is not well-controlled. This combination of traits indicates dysfunction of the neuroendocrine system. In PWS patients, the paternal copy of PWS genes, situated on chromosome 15q11-q13, is missing or silenced. This region includes *SNORD116* and *SNORD115* snoRNA clusters, *SNRPN, SNURF, NDN, MKRN3, and MAGEL2*. These genes are present on the maternal copy of chromosome 15 but silenced by hypermethylation and repressive histone modulation of the PWS imprinting center (PWS-IC) (Cassidy et al., 2000; Fulmer-Smentek and Francke, 2001; Henckel et al., 2009). Genetic ablation studies in murine models attempting to dissect the genetic underpinnings of PWS have converged on the *SNORD116* snoRNA cluster as the major contributor to PWS etiology (Bortolin-Cavaille and Cavaille, 2012; Burnett et al., 2017a; Ding et al., 2008; Gallagher et al., 2002; Polex-Wolf et al., 2018; Qi et al., 2016; Zhang et al., 2012). However, human genetic and clinical data suggested that other PWS genes, such as *NDN, MAGEL2* and *SNRPN* also contribute to PWS etiology by regulation of hypothalamic neural function (Kuslich et al., 1999; Matarazzo et al., 2017; Miller et al., 2009; Wijesuriya et al., 2017).

There is currently no cure for PWS, with most clinical efforts focusing on hormonal management to modulate the neuroendocrinal circuit (Angulo et al., 2015; Dykens et al., 2018; Einfeld et al., 2014; Miller et al., 2017; Tauber et al., 2017). However, owing to relatively clear genetic root cause for PWS, epigenetic activation of the transcriptionally silent maternal PWS genes represents an attractive therapeutic approach. Indeed, 5-Azacytidine (5-Aza), a cytidine analog that antagonizes *de novo* DNA methylation has been proven to demethylate PWS-IC and substantially reactivate maternal PWS genes in mitotic cells (Saitoh and Wada, 2000; Takano et al., 2007). In addition, deregulation of the repressive histone methyltransferase *SETDB1* by genetic ablation (Cruvinel et al., 2014), and application of HDAC inhibitors (Saitoh and Wada, 2000) have demonstrated variable effects on PWS gene derepression in mouse models and various human cells. A recent report demonstrated that inhibition of the euchromatin histone methyltransferase G9a by small molecules could reactivate *SNORD116* and other PWS genes in PWS patient fibroblasts, consistent with the key role that G9a plays in catalyzing H3K9 dimethylation at the maternal PWS-IC to silence PWS genes. Notably, G9a inhibition also significantly improves the life span of a PWS mouse model, which otherwise will die within 2 weeks after birth (Kim et al., 2017). Interestingly, both G9a and *SETDB1* are histone methyltransferases that methylate H3K9 and have been demonstrated to form a megacomplex that interacts with *de novo* and maintenance DNA methyltransferases (DNMTs) and thus is linked to PWS-IC methylation and genetic imprinting in rodent development and human iPSCs, although the detailed molecular mechanism remains elusive (Fulmer-Smentek and Francke, 2001; Henckel et al., 2009; Xin et al., 2003; Zhang et al., 2016). These findings support the notion that there is a molecular interplay between DNA methylation and heterochromatic modulation of the PWS-IC in maintaining genomic imprinting at the PWS locus.

Modeling genetic disease *in vitro* using patient derived iPSCs has been a powerful approach to study disease mechanisms (Soldner and Jaenisch, 2018). A body of work has been reported on *in vitro* disease modeling for PWS, not only providing valuable patient-derived, genetically accurate disease cell lines for the PWS community, but also elucidating new mechanisms that have valuable clinical implications (Burnett et al., 2016; Chamberlain et al., 2010; Kim et al., 2013; Martins-Taylor et al., 2014; Polvora-Brandao et al., 2018; Takano et al., 2007; Yang et al., 2010). In the present study, we model PWS using patient-specific iPSC-derived neural progenitor cells (NPCs) and cortical excitatory neurons, and quantitatively profile the expression of PWS genes using a NanoString panel. We demonstrate that methylation of the PWS-IC is stable during neuronal lineage conversion, and that the maternal PWS genes remain silenced in PWS NPCs and neurons. We show that multiple small molecule inhibitors of G9a activate maternal PWS genes in a dose dependent manner in both NPCs and neurons via PWS-IC methylation independent mechanisms. In addition, we found that G9a inhibitors in PWS neurons induce *GNRH1* and *HTR2C*, two neuron specific genes associated with PWS pathology (Miller et al., 2009; Glatt-Deeley et al., 2010; Garfield et al., 2016). Our data suggest that deregulation of the H3K9 methyltransferase G9a can partially activate maternal PWS genes in neural progenitors and neurons, the cell types of direct pathophysiological relevance for PWS. We therefore provide proof of concept for activation of critical PWS genes *via* epigenetic modulation.

## Results

### Modeling PWS disease in a dish using patient derived neural progenitors and neurons

To test the idea of reactivation of PWS genes in patient derived neural cells, we sought to generate neural progenitor cells (NPCs) and neurons from a PWS patient iPSCs (PWS1-7) who has a ∼ 5 Mb deletion from BP2-BP3 along the paternal PWS locus, spanning from *MKRN3* at the centromeric end to *HERC2* at the telomeric end (Figure 1A). MCH2-10 was included in the study as a healthy control iPSC line (Chamberlain et al., 2010; Cruvinel et al., 2014). NPCs were derived from these two iPSC lines using a dual SMAD inhibition approach, and high purity excitatory neurons were derived from NPCs using a doxycycline inducible NGN2 transgene (Figure 1B). Immunostaining analyses of NPCs and neurons suggested robust neural cell differentiation from patient derived iPSCs. NPCs expressed *SOX2* and PAX6, two major neural progenitor markers and NGN2-iduced neurons expressed neuron specific markers TUJ1 and MAP2 without astroglia contamination as indicated by the negative staining of glia specific marker GFAP. No differentiation deficits were observed for PWS patient derived iPSCs compared to WT iPSC (Figure 1C&D). Gene profiling analysis in Figure 1E suggest that NPCs from both WT and PWS iPSCs express high level of neural progenitor specific genes *SOX2* and low level of neuron specific genes. In contrast, neurons from both WT and PWS iPSC lines expressed low levels of *SOX2*, but high levels of neural differentiation genes *DCX* and *NEUROD2*, the pan neuronal gene *TUBB3*, and neural maturation genes *RBFOX3* and *SYN1*. In addition, no dividing cells were detected in NGN2-induced neurons over the course of the neuronal culture (data not shown).

**Figure 1.**
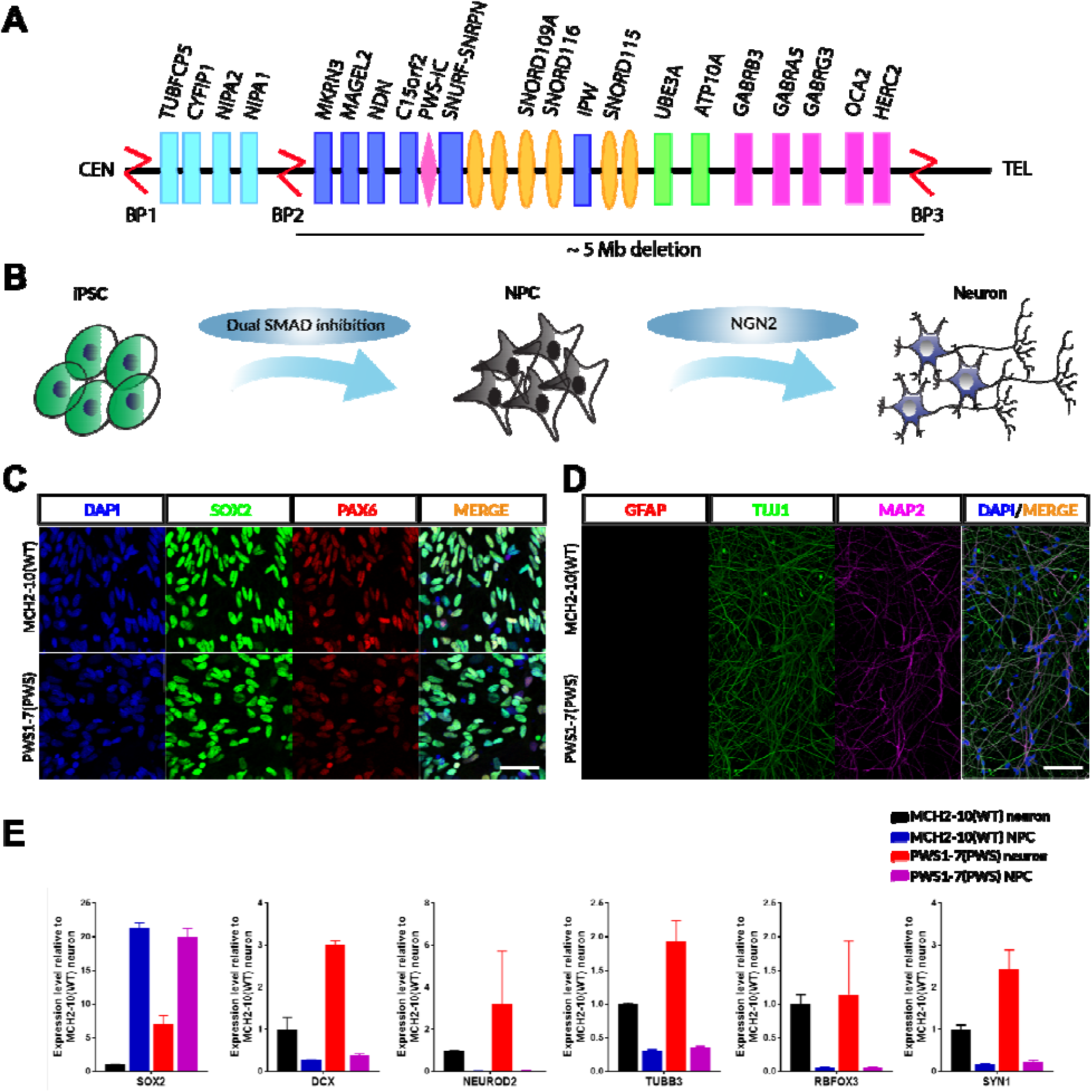
Modeling PWS in a dish using patient derived NPCs and Neurons. (A). Schematic diagram depicting the genome organization of part of chromosome 15q11-q13 and canonical breakpoint (BP) deletions found on the paternal alleles in PWS patients. A subset of coding and non-coding genes are labeled on the locus. PWS1-7 patient iPSCs used in this study harbor a ∼ 5 Mb deletion (BP2-BP3) that spans from *MKRN3* to *HERC2* (centromeric to telomeric). (B). Schematic diagram of the cortical neuron differentiation process. (C&D). Representative confocal images of differentiated NPCs and neurons characterized by the following markers: *SOX2* (green) and PAX6 (red), markers for NPCs; TUJ1 (green) and MAP2 (magenta), specific markers for neuronal axons and dendrites; GFAP (red), astroglia marker which is negative in differentiated WT and PWS neurons. Cells were counterstained with DAPI (blue). Scale bar: 100 um. (E). Expression profile of a set of neuronal differentiation genes by NanoString analysis in differentiated WT and PWS NPCs and neurons: *SOX2*, early neuronal progenitor gene; *DCX* and *NEUROD2*, early neuronal differentiation genes; *TUBB3*, neurite specific gene; *RBFOX3* and *SYN1*, neural maturation genes. Data are normalized to gene expression of MCH2-10 WT NPCs. Bars represent mean +/− SD of two biological replicates.

### Methylation of PWS-IC does not change during neuronal differentiation in vitro

To test if differentiated PWS NPCs and neurons serve as robust in vitro PWS models, we initially analyzed the epigenetic stability of the PWS-IC by measuring the methylation of 8 CpG dinucleotides within the PWS-IC near exon 1 of the *SNRPN* gene in PWS NPCs and neurons (White et al., 2006). Pyrosequencing results demonstrated that the PWS-IC remains hypermethylated in PWS NPCs and neurons (70%-100% methylation) and hemimethylated (∼ 50% methylation) in WT NPCs and neurons, suggesting that the PWS-IC does not change during neuronal differentiation in vitro (Figure 2A). To determine whether appropriate PWS-relevant gene expression was also maintained during neuronal differentiation, we designed a NanoString panel to quantitatively measure the expression of paternally-expressed, imprinted genes in the PWS region. This panel included snoRNA clusters *SNORD116, SNORD115, SNORD64* and SNORD109, as well as *IPW, MAGEL2, MKRN3, NDN, SNRPN* and *SNURF* (Table S3). All of the paternally-expressed imprinted genes in the PWS region remain repressed in PWS NPCs and neurons, indicating that the genetic imprinting of the PWS locus is not altered during neuronal differentiation. Interestingly, the relative expression of these transcripts in PWS cells relative to WT cells is different between NPCs and neurons, suggesting possible cell type specific regulation of the genes within the PWS locus (Figure 2B&C).

**Figure 2.**
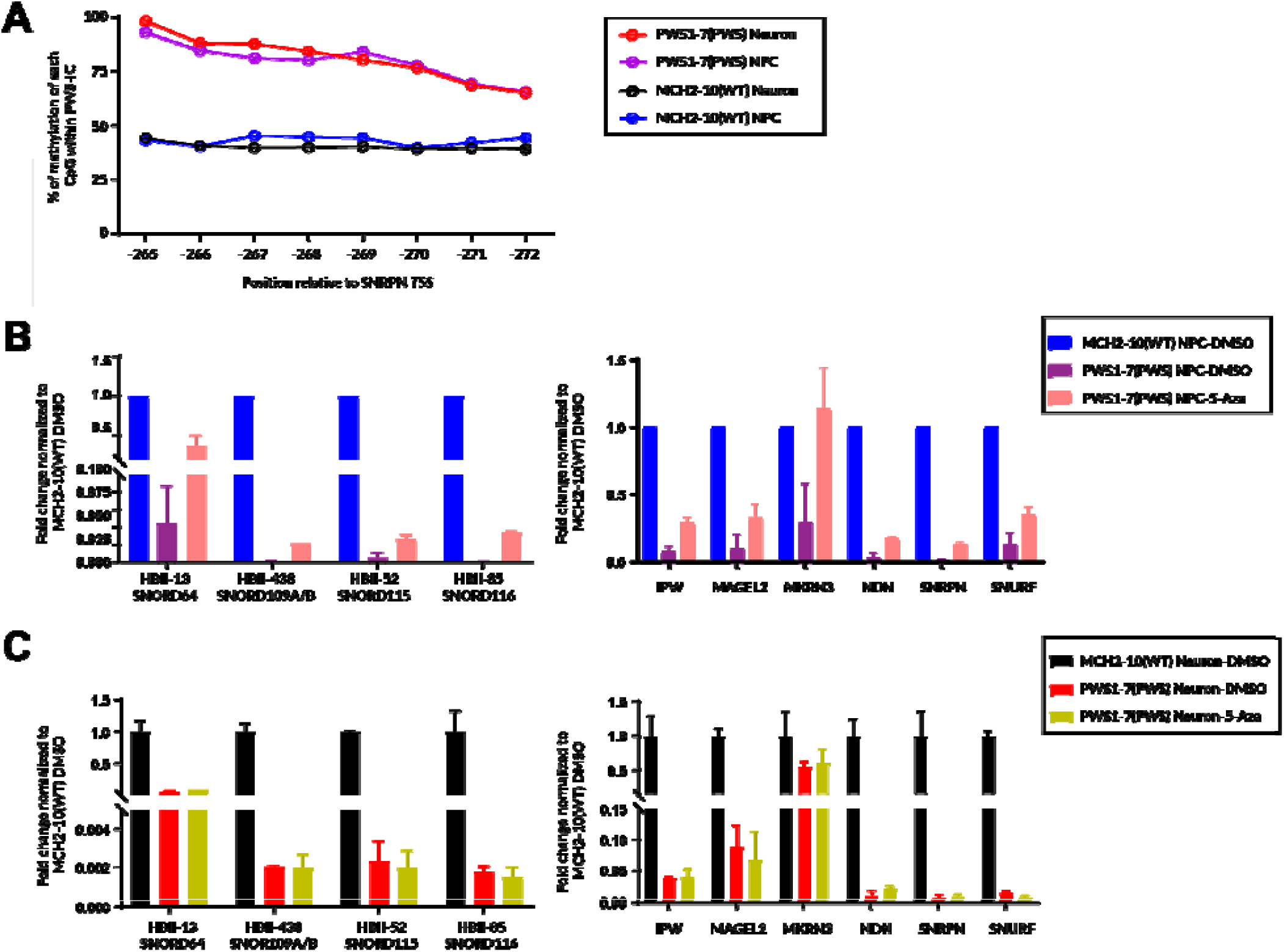
5-Aza reactivates PWS gene expression in PWS patient NPCs but not neurons. (A). Pyrosequencing results demonstrating that methylation of the PWS-IC is refractory to neuronal differentiation in vitro. The PWS-IC remains hypermethylated in PWS NPCs and neurons (> 80% methylation on average across the 8 CpG methylation sites) and hemimethylated (∼ 50% methylation on average across the 8 CpG methylation sites) in WT NPCs and neurons. (B). PWS genes, including *SNORD116*, remain silenced in PWS NPCs. Many PWS genes are significantly activated to greater than 20% of the MCH2-10 WT NPCs in response to 1 µM 5-Aza treatment for one week. (C). PWS genes remain silenced in PWS neurons, and 5-Aza treatment for two weeks does not affect PWS gene expression in PWS neurons. Results are compared to gene expression levels of the MCH2-10 WT NPCs and neurons. Bars represent mean +/− SD of two biological replicates.

### 5-Aza reactivates PWS gene expression in PWS patient NPCs but not neurons

5-Azacytidine (5-Aza), the cytidine analog known to block de novo DNA methylation, has been used to activate genes whose promoters are hypermethylated (Bar-Nur et al., 2012; Saitoh and Wada, 2000; Takano et al., 2007). As all the previous attempts to activate epigenetically silenced genes were demonstrated in mitotic cells or heterogenous culture with mitotic cell contamination (e.g. lymphoblastoids and fibroblasts), we asked if 5-Aza could similarly impact PWS gene expression in postmitotic neuronal cells. Figure 2B&C demonstrate that 5-Aza treatment significantly activates many PWS genes to greater than 20% of the WT in PWS NPCs but shows no effect in PWS neurons. These results suggest a lack of the antagonizing effect of 5-Aza on DNA methylation in postmitotic neurons. This is consistent with the fact that there is no prominent de novo DNA methylation, which requires DNA replication, in neurons (Lister and Mukamel, 2015).

### G9a inhibition activates PWS genes in patient-derived NPCs

The G9a histone H3K9 methyltransferase (HMT) was shown to be required for PWS-IC methylation maintenance in rodents (Henckel et al., 2009; Xin et al., 2003), and inhibition of G9a by small molecules has been reported to reactivate maternal *SNORD116* in PWS fibroblasts and in a *Snord116*^*p-/m+*^ mouse model (Kim et al., 2017). To examine the effect of G9a inhibition on maternal PWS gene activation, we treated the PWS NPCs with well-characterized G9a small molecule inhibitors with similar potency and selectivity (A-366, UNC-0638 and UNC-0642 (Kubicek et al., 2007; Kim et al., 2017)) for one week and observed *SNORD116* gene reactivation in PWS NPCs in a dose dependent manner from 1 µM to 10 µM (Figure 3A). Among the three compounds, UNC-0638 demonstrated slightly better efficacy in neural progenitors, with the 10 µM treatment leading to higher levels of *SNORD116* expression than 1 µM 5-Aza treatment (Figure 3A). For subsequent studies, we used UNC-0638 as the tool compound for G9a inhibition. Gene profiling of the entire PWS gene panel suggested that inhibition of G9a by UNC-0638 activated other snoRNA clusters (*SNORD115, SNORD64* and *SNORD109A/B*) and PWS genes (*MKRN3*, MAGEL2, *SNRPN, SNURF, IPW* and *NDN*) in PWS NPCs in a dose dependent manner. Greater activation was consistently seen across the maternal PWS genes in PWS NPCs treated with 10 µM UNC-0638 than 1 µM 5-Aza (Figure 3B).

**Figure 3.**
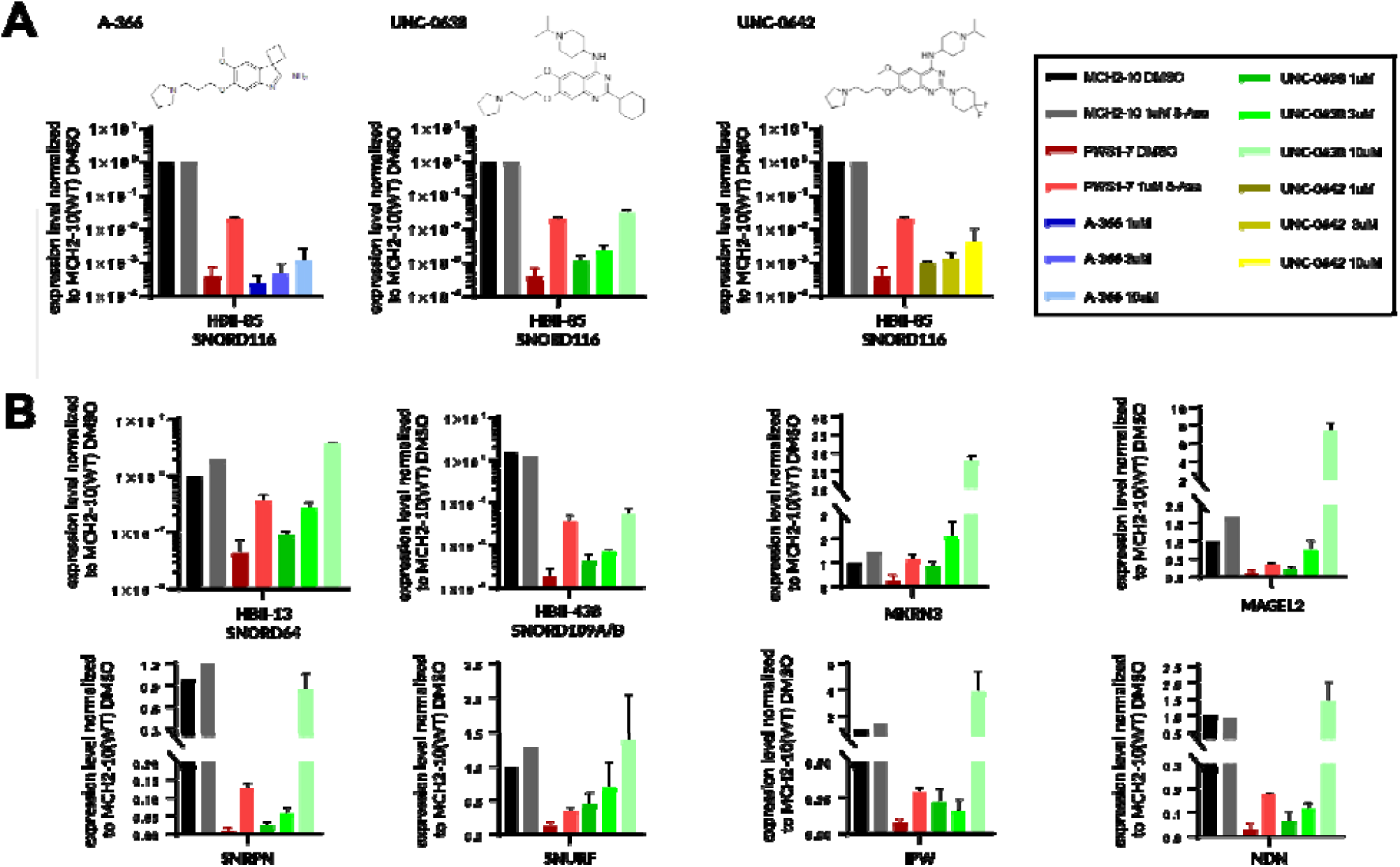
G9a inhibition activates PWS genes in patient derived NPCs. (A). Multiple G9a inhibitors (A-366, UNC-0638 and UNC-0642) activate *SNORD116* gene expression in PWS NPCs in a dose dependent manner (1 µM, 3 µM and 10 µM). (B). Inhibition of G9a by UNC-0638 activates other snoRNA genes *SNORD64* (HBII-13) and *SNORD109A/B* (HBII-438), and a panel of PWS genes (*MKRN3*, MAGEL2, *SNRPN, SNURF, IPW* and *NDN*) in PWS NPCs in a dose dependent manner. Results are compared to 1 µM 5-Aza treatment and WT NPCs. NPCs were treated with DMSO or compounds for one week. Data are normalized to expression of genes in MCH2-10 NPCs with DMSO treatment. Bars represent mean +/− SD of two biological replicates.

### G9A inhibition activates PWS genes in patient-derived neurons

The EC_50_ of UNC-0638 in mouse fibroblast cells was reported at ∼ 1.6 µM (Kim et al., 2017), and we observed cellular toxicity when 10 µM UNC-0638 was used for neuronal treatment (data not shown). We therefore decided to treat the PWS patient derived neurons with UNC-0638 at 3 µM for 2 weeks. Gene profiling using the NanoString panel suggested that inhibition of G9a activates snoRNAs (*SNORD115, SNORD116, SNORD64* and *SNORD109A/B*, left panel) and other PWS genes (*MKRN3*, MAGEL2, *SNRPN, SNURF, IPW* and *NDN*, right panel) at various levels in PWS neurons. *SNORD116* was only induced to ∼ 1% of the WT neurons, and MAGEL2 and some other PWS genes were induced to levels equal to or greater than WT levels (Figure 4A).

**Figure 4.**
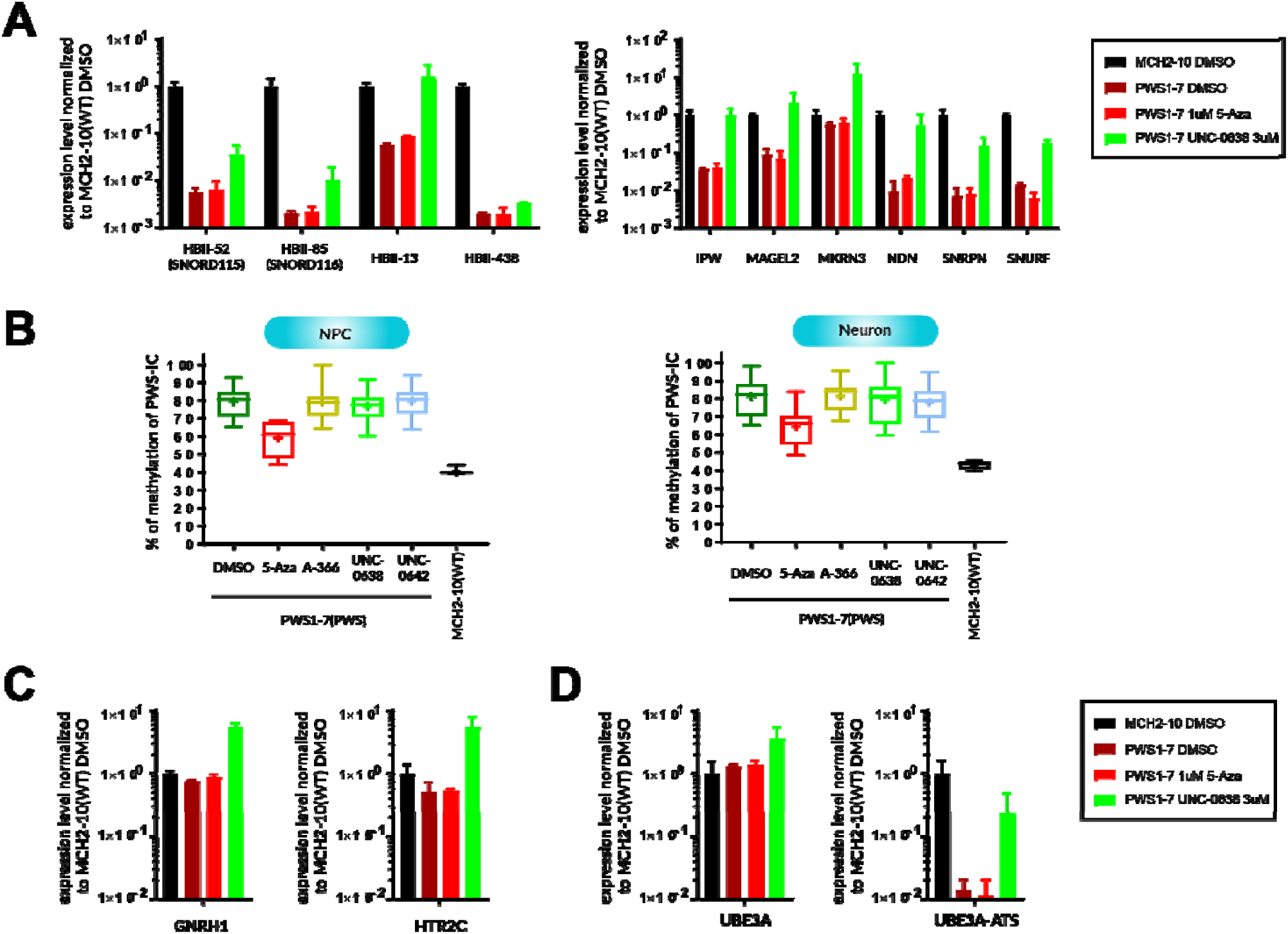
G9a inhibition activates *SNORD116* and PWS genes in patient neurons via IC-methylation independent mechanism. (A). Inhibition of G9a by UNC-0638 at 3 µM activates snoRNAs (*SNORD115, SNORD116, SNORD64* and *SNORD109A/B*, left panel) and other PWS genes (*MKRN3*, MAGEL2, *SNRPN, SNURF, IPW* and *NDN*, right panel) in PWS neurons. The level of gene activation varies, with the *SNORD116* only induced to ∼ 1% of the WT neurons, and MAGEL2 and other PWS genes to almost 100% of the WT neurons. (B). Pyrosequencing results demonstrating that whereas 1 µM 5-Aza reduces PWS-IC methylation in PWS1-7 patient NPCs, G9a inhibition does not affect methylation of the PWS-IC in both PWS NPCs and neurons. Results are compared to WT and DMSO treated PWS NPCs and neurons. Data were plotted as box and whisker plots with a “+” denoting the mean of all 8 CpG methylation sites and the box indicating 25 and 75 percentiles. (C). Inhibition of G9a induces expression of reported neuronal specific PWS downstream genes *GNRH1* and *HTR2C*. (D). Inhibition of G9a induces expression of *UBE3A* and *UBE3A-ATS* genes adjacent to the PWS locus. Data are normalized to expression of genes in MCH2-10 neurons with DMSO treatment. Bars represent mean +/− SD of two biological replicates. NPCs and neurons were treated with DMSO or compounds for one week and two weeks, respectively.

### G9-mediated activation occurs via a PWS-IC-methylation independent mechanism

As G9a has been linked to the maintenance of methylation at the PWS-IC, and 5-Aza mediated DNA demethylation leads to partial activation of the maternal PWS genes, we sought to investigate if G9a inhibition induced PWS gene activation by causing PWS-IC demethylation. Pyrosequencing results demonstrate that whereas 1 µM 5-Aza reduces PWS-IC methylation in PWS1-7 patient NPCs, G9a inhibition does not affect methylation of the PWS-IC in both PWS NPCs and neurons (Figure 4B), indicating that G9a inhibition-induced PWS gene activation in patient NPCs and neurons likely occurs through an PWS-IC methylation-independent mechanism. This data is consistent with the report by Kim et al in PWS rodent model and patient fibroblasts (Kim et al., 2017).

### Inhibition of G9a also affects *UBE3A* related genes and PWS downstream genes in PWS neurons

We finally examined two non-15q genes that are reported to be affected in PWS patient neurons, presumably downstream of the PWS-causing mutation (Miller et al., 2009; Glatt-Deeley et al., 2010; Garfield et al., 2016). Figure 4C shows that inhibition of G9a induces expression of the neuron specific downstream genes *GNRH1* and *HTR2C* by 6-8-fold (Figure 4C). In addition, *UBE3A* and *UBE3A-ATS* genes are also induced significantly by G9a inhibition in PWS patient derived neurons (Figure 4D). The functional consequence of the upregulation of these genes in PWS neurons remains to be further investigated.

## Discussion

Modeling imprinting disorders such as PWS using animal models remains challenging. For example, murine models usually do not recapitulate the diverse segmental deletion haplotypes on the 15q11-q13 in PWS patients, and do not recapitulate the whole array of PWS patient phenotypes, including obesity, a phenotype central to the disorder (Burnett et al., 2017a; Ding et al., 2008; Garfield et al., 2016; Matarazzo et al., 2017; Qi et al., 2016). In addition, lengthy transgenerational mouse crossing is required to precisely capture the parent-of-origin imprinting genetics of PWS. As such, patient derived cells represent a complementary, yet more disease relevant model for PWS. Indeed, new mechanisms that show valuable clinical implications for PWS have been elucidated using diseased cells with direct pathophysiological relevance (Burnett et al., 2016; Chamberlain et al., 2010; Polvora-Brandao et al., 2018; Yang et al., 2010). For instance, Cruvinel et al. reported that knocking down the H3K9 methyltransferase *SETDB1* activates maternal PWS genes in patient derived iPSCs (Cruvinel et al., 2014). Following that, Langouët et al. observed partial restoration of maternal PWS genes in PWS neurons by ablating ZNF274, a KRAB-domain zinc finger protein purported to interact with *SETDB1* (Langouet et al., 2018). More recently, human hypothalamic-like neurons have been differentiated from iPSCs and used to model PWS. Prohormone convertase PCSK1 deficiency has been identified as one a putative major contributor to the neuroendocrine phenotype of PWS. This result was further supported by the transcriptional profiling of hypothalami from the *Snord116*^*p-/m+*^ mouse model (Bochukova et al., 2018; Burnett et al., 2017b; Wang et al., 2015; Zhang et al., 2012). In addition, *SNORD116* has been indicated to play the most critical role in PWS etiology, which is corroborated with various genetic ablation studies using murine models (Bortolin-Cavaille and Cavaille, 2012; Burnett et al., 2017a; Ding et al., 2008; Gallagher et al., 2002; Polex-Wolf et al., 2018; Qi et al., 2016; Zhang et al., 2012). In this study, we derived neural progenitor cells and NGN2-induced glutamatergic neurons with high purity from PWS patient iPSCs with a ∼ 5 Mb deletion on the paternal PWS locus (PWS1-7) (Figure 1A&B). We demonstrated that the methylation of the PWS-IC does not change during directed neuronal lineage conversion, and that maternal *SNORD116, NDN*, MAGEL2 and *SNRPN*, the key genes responsible for PWS etiology, remain silenced in differentiated PWS NPCs and neurons (Figure 2A-C). Our characterization of PWS1-7 neural cells suggest that NPCs and neurons are suitable in vitro disease models with satisfying genetic and epigenetic stability.

We found that 1 µM 5-Aza activates expression of many maternal PWS genes to ∼20%-100% of levels seen in NPCs generated from a WT control line. This result confirmed the previous findings that deregulating PWS-IC methylation by 5-Aza is able to activate the maternal alleles of PWS genes in mitotic cells (Saitoh and Wada, 2000; Takano et al., 2007). However, our data also showed that 5-Aza does not cause PWS gene derepression in neurons. This may be due to the lack of de novo DNA methylation machinery in postmitotic cells such as neurons (Figure 2).

Using multiple selective compounds with similar potency (IC_50_ at nanomolar scale for A-366, UNC-0638 and UNC-0642), we found that G9a inhibition activates maternal PWS genes in a dose dependent manner in both NPCs and neurons (Figure 3 and 4A). Interestingly, G9a inhibition does not induce methylation changes of the PWS-IC, indicating that reduction of H3K9 dimethylation alone suffices to induce an open chromatin state within the PWS locus that leads to partial activation of the maternal PWS genes (Figure 4B). It is noteworthy that *SNORD116*, the gene likely responsible for most of the PWS phenotypes, is only activated to ∼ 1% of levels seen in neurons from a healthy individual. The functional consequence of such a low gene dosage for *SNORD116* needs to be further investigated in PWS neurons.

Genomic analyses in mouse and human have identified potential PWS downstream genes that show etiological contributions to the neuroendocrinal deficits in PWS (Bochukova et al., 2018; Burnett et al., 2017b; Galiveti et al., 2014; Zhang et al., 2012). Among them, the gonadotropin-releasing hormone 1 (*GNRH1*) has been shown to be downregulated by *NDN* deficiency in murine hypothalamic cells (Miller et al., 2009). Similarly *HTR2C*, the serotonin receptor gene mediating appetite control, has been reported to be mis-expressed due to paternal *SNORD115* deficiency in both PWS cells and a rodent model (Garfield et al., 2016; Glatt-Deeley et al., 2010; Kishore and Stamm, 2006; Morabito et al., 2010). Interestingly, our data showed that both genes were significantly induced by G9a inhibition (Figure 4C). Whether these genes are directly induced by G9a inhibition or secondary to the partial reactivation of PWS genes remains to be further investigated. In addition, the functional consequence of G9a inhibitor-induced partial activation of PWS genes and induction of *GNRH1* and *HTR2C* in neurons, especially hypothalamic-like neurons, remains an interesting topic to study in the future. Of note, while Kim et al. showed no overtly altered expression of Ube3a and Ube3a-ATS following G9a inhibition in a PWS mouse model (Kim et al., 2017), our human neuron data suggested that both *UBE3A* and *UBE3A-ATS* expression are significantly increased by G9a inhibition (Figure 4D). Increased *UBE3A-ATS* expression is consistent with activation of *SNORD116* and *SNORD115*, since they are part of the same transcriptional unit. However, *UBE3A-ATS* may not be sufficiently upregulated to repress *UBE3A* (Hsiao et al., 2019; Langouet et al., 2018). Although it is not clear why *UBE3A* expression is increased following G9a inhibition, this result may alleviate one potential safety concern related to gene activation specificity at this locus. Further studies are necessary to fully address this concern. In summary, using disease relevant cellular models, our study provided in vitro proof of principle for using epigenetic intervention as a potential therapeutic approach to PWS treatment.

## Materials and Methods

### iPSC culture

WT control (MCH2-10) and PWS (PWS1-7) iPSC lines used in this study are listed in Table S1. iPSCs were cultured in Geltrex (ThermoFisher Scientific, A1413302) coated flasks in StemFlex medium (ThermoFisher Scientific, A3349401) and fed with fresh media every other day. Cells were passaged using ReLeSR™ (STEMCELL Technology, 05873) and subjected to mycoplasma and karyotypic analysis over the passages to ensure culture sterility and genome stability. iPSCs were cryopreserved in CryoStem™ hPSC Freezing Medium (Biological industries, 05-710-1E). To recover iPSCs, vials were thawed with gentle swirling in a 37°C water bath for 1-3 min until no ice piece was observed in the vial. Cells were immediately transferred into a 50 ml tube. 5 ml StemFlex™ media were added dropwise into the cells slowly. Cells were then spun at 200 × g for 5 min at room temperature, and resuspended in 5 ml StemFlex™ media with 1 × RevitaCell supplement (ThermoFisher Scientific, A2644501) for plating for the first 24 hrs.

**Table S1:**
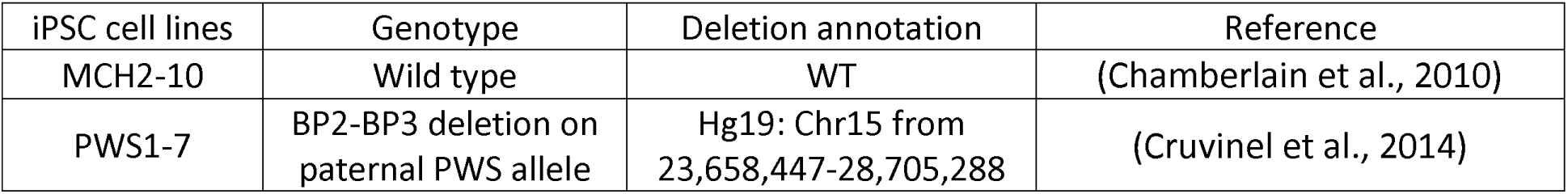
iPSC lines used in this study

### NPC generation

MCH2-10 WT and PWS1-7 neural progenitor cells were generated with dual SMAD inhibition approach using PSC Neural Induction medium (ThermoFisher Scientific, A1647801) according to manufacturer’s instructions. Briefly, iPSCs were cultured on StemFlex™ medium in Geltrex coated flasks, when reaching ∼80% confluency, iPSCs were dissociated with pre-warmed Accutase™ Cell Dissociation Reagent (ThermoFisher Scientific, A1110501) at 37°C for 5 min. Cell pellets were resuspended in 5 ml StemFlex medium after centrifugation at 200 × g for 5 min. Cell density was calculated using Countess II FL Automated Cell Counter (ThermoFisher Scientific). To induce the NPC differentiation, iPSC single cells were plated in Geltrex coated 6-well plate at 2.5 × 10^5^ /cm^2^ in StemFlex medium with 1 × RevitaCell supplement. On day two post plating, feed the cells with Neural Induction medium (Neurobasal medium + 1 × neural induction supplement). Feed the cells everyday with 5 ml Neural Induction medium for 5 days. Neural progenitor cells were then cryopreserved at P0 using STEMdiff™ Neural Progenitor Freezing Medium (STEMCELL Technologies, 05838) and further expanded with neural progenitor cell expansion medium (50% Neurobasal medium + 50% Advanced DMEM/F-12 medium + 1 × neural induction supplement). Usually when plated at 2 × 10^6^/well, cells will reach 100% confluence in 4-5 days in 6-well plate. NPCs were then cultured in STEMdiff™ Neural Progenitor Medium (STEMCELL Technologies, 05833) for maintenance, neuronal differentiation and compound treatment. NPCs maintain their cellular fate up to 8-10 passages in NPC medium.

### Neuronal differentiation from NPCs

Cortical excitatory neurons were derived from MCH2-10 and PWS1-7 NPCs using a Dox inducible NGN2 transgene according to a modified method from Zhang et al. (Zhang et al., 2013). Briefly, on day 0, WT and PWS NPCs cultured on Geltrex coated flasks in STEMdiff™ Neural Progenitor Medium were dissociated with StemPro™ Accutase™ Cell Dissociation Reagent as single cells. Cells were infected with pTet-O-NGN2-2A-Puro virus (1 × 10^9^ IFU/ml) in suspension at MOI of 1. Infected cells were cultured on Geltrex in NPC medium with 1 × RevitaCell supplement and 2 ug/ml Doxycycline (Clontech) to induce NGN2 gene expression. To select the infected cells, cells were fed with NPC medium with 2 ug/ml Doxycycline and 3.34 ug/ml Puromycin (ThermoFisher Scientific). On Day 3, the cells were dissociated with pre-warmed Accutase and re-plated as single cells into PDL and 50 µg/ml Laminin (Sigma-Aldrich, L2020) coated plate with appropriate density in NBM medium (Neurobasal™ Media with 1 × NEAA (ThermoFisher Scientific), 1 × Glutamax, 0.3% D-(+)-Glucose, 2 ug/ml Doxycycline and 3.34 ug/ml Puromycin, 1 × N27 supplement, and 10 ng/ml BDNF (STEMCELL, 78005) and GDNF (R&D Systems, 212-GD)). Replace half of the media with fresh NBM with 1 × penicillin/streptomycin every 7 days before the cells were collected for downstream analysis.

### Compound treatment

WT and PWS NPCs and neurons were cultured in 96 well plates at 100,000 cells per well for compound treatment. DMSO, the compound solvent, was used as treatment control for various concentrations of 5-Aza and G9a inhibitors A-366, UNC-0638 and UNC-0642. DMSO concentration was fixed at 0.1% for all the treatments. NPCs were treated with compounds for one week whereas neurons were treated for two weeks with retreatment each week.

### Plasmid and Lentivirus

The lentiviral packaging vector pTetO-NGN2-T2A-Puro (Addgene Plasmid#52047) (Zhang et al., 2013) was a generous gift from Dr. Marius Wernig at the Institute for Stem Cell Biology and Regenerative Medicine, Stanford University. The plasmid was maxi-prepared, and sequence confirmed before lentiviral packaging and production at 1 × 10 _9_ IFU/ml by Alstem LLC.

### Immunofluorescence and imaging analysis

NPCs and neurons were fixed with 4% paraformaldehyde (PFA) for 10 min at room temperature. Cells were permeabilized with PBST (1 × PBS solution with 0.5% Triton X-100) before blocking with 5% BSA in PBST. Cells were then incubated with appropriately diluted primary antibodies in PBST with 1% BSA for 1 hours at room temperature or overnight at 4°C. After washing with PBST for 3 × 10 min at room temperature, cells were incubated with fluorophore conjugated secondary antibodies and DAPI. NPCs and neurons were washed 3 × 10 min with PBST at room temperature before being subjected to imaging using LSM700 confocal microscopy (ZEISS). Data were processed with an imaging analysis freeware FIJI (ImageJ, https://fiji.sc/). Antibodies used in this study are listed in Table S2.

**Table S2:** Antibodies and dilution for NPC and neuron characterization

| Antibodies | Vendor | Cat. No. | Dilution |
| --- | --- | --- | --- |
| Rabbit anti PAX6 | BioLegend | 901301 | 1:200 |
| Rabbit anti GFAP | Dako | Z0334 | 1:250 |
| Mouse anti MAP2 | ThermoFisher Scientific | MA5-12826 | 1:150 |
| Goat anti SOX2 | R&D Systems | AF2018 | 1:150 |
| Chicken anti TUJ1 | AVES | TUJ89947984 | 1:500 |
| DAPI | Sigma-Aldrich | 10236276001 | 1:10,000 |
| Alexa Fluor 647nm Donkey anti-Mouse IgG (H+L) | Invitrogen | A-31571 | 1:2,000 |
| Alexa Fluor 488 nm Goat anti-Chicken IgY (H+L) | Invitrogen | A-11039 | 1:2,000 |
| Alexa Fluor 555 nm Donkey anti-Rabbit IgG (H+L) | Invitrogen | A-31572 | 1:2,000 |
| Alexa Fluor 488 nm Donkey anti-Goat IgG (H+L) | Invitrogen | A-11055 | 1:2,000 |

### NanoString probe design and data analysis

NanoString probe panels for gene expression profiling were designed by NanoString Technologies LLC (Table S3 and S4). Hybridizing probes were synthesized by Integrated DNA Technologies (IDT). Total RNAs were prepared in RLT RNA lysis buffer supplemented with 1:100 β-mercaptoethanol (Qiagen) at 5,000 cells/µl. Cells were lysed at room temperature with agitation for 10 min and RNA lysates were kept at −80°C before being subjected to gene profiling analysis. 2 µl RNA lysates were loaded into the NanoString nCounter^®^ cartridge for gene expression assay. Data were analyzed using nSover3.0 software and plotted using Prism Graphpad. Gene expression levels were normalized to the housekeeping gene GAPDH for PWS gene profiling (Table S3) and the average expression of the 4 housekeeping genes PUM1, PPIA, PSMC4 and ACTB for NPC and neuron QC (Table S4). Statistics were calculated with unpaired t-Test.

**Table S3:**
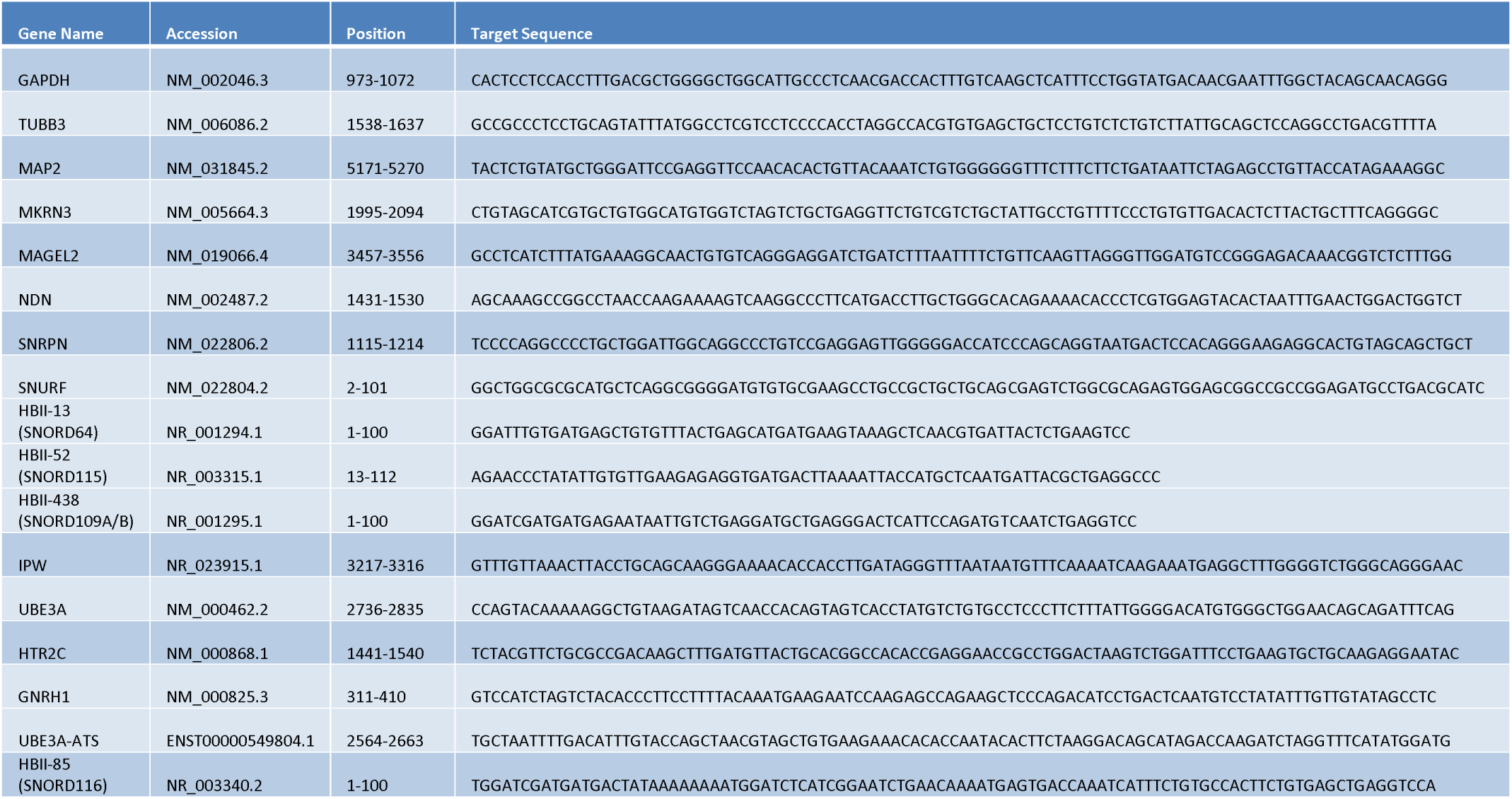
NanoString panel for expression profiling of PWS locus and PWS responsive genes

**Table S4:**
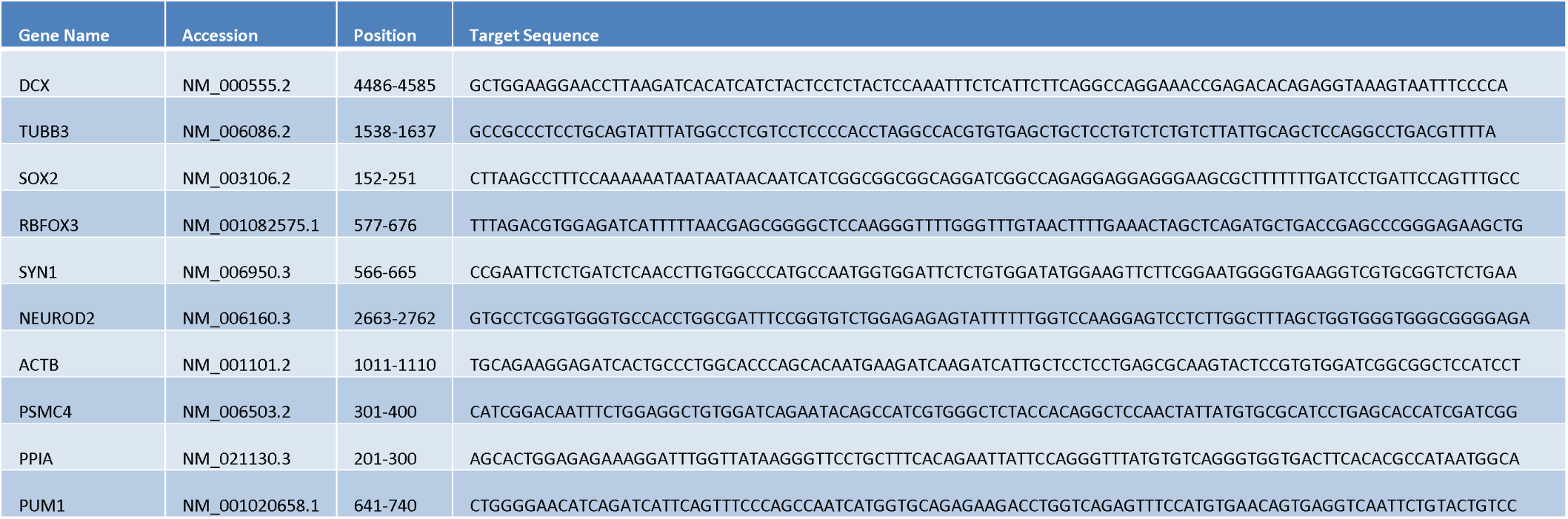
NanoString gene panel for characterizing NPCs and neurons

### PWS-IC methylation analysis

PWS-IC methylation was analyzed by pyrosequencing of the intron 3 in human *SNRPN* gene (EpigenDX, Assay ID: ADS265-RS1), an established assay to determine the methylation status of PWS-IC (White et al., 2006). The CpG island in the intron 3 of the *SNRPN* gene contains 8 CpG dinucleotide methylation sites. To perform the pyrosequencing, genomic DNA from compound treated NPCs or neurons were extracted using DNA Lysis buffer (100 mM Tris-Cl, 50 mM EDTA, 1% (w/v) SDS and 50 µg/ml Proteinase K, pH 8.0), subsequently precipitated with 1:1 isopropanol and purified with 70% EtOH. DNA was eluted in TE buffer (1 mM Tris and 1 mM EDTA, pH 8.0) and incubated at 50°C to ensure complete dissolution in TE buffer. DNA was bioanalyzed for genomic integrity, concentration and purity before being subjected to the pyrosequencing for methylation assay.

## Author Contributions

H.W., and A.C. conceived the idea for this project. H.W. designed the experiments and interpreted the data. H.W., C.N., V.V. performed the experiments. S.C. provided MCH2-10 WT and PWS1-7 PWS patient iPSC lines. H.W. wrote the manuscript with input from all the other authors.

### ACKNOWLEDGMENTS

We thank Dr. Stormy Chamberlain at Genetics and Genome Sciences, University of Connecticut Health Center for key PWS iPSC lines used in this study. We thank Ms. Jennifer Love at Whitehead genomic center for NanoString gene expression profiling analysis, Dr. Marius Wernig for pTetO-NGN2-T2A-Puro plasmid, and the bioinformatic group at NanoString Technologies Inc. for designing the Neuronal QC and PWS NanoString probe panels. We thank members of Fulcrum Therapeutics for discussions and suggestions on the manuscript.

